# A programmable pAgo nuclease with universal guide and target specificity from the mesophilic bacterium *Kurthia massiliensis*

**DOI:** 10.1101/2021.02.03.429301

**Authors:** Ekaterina Kropocheva, Anton Kuzmenko, Alexei A. Aravin, Daria Esyunina, Andrey Kulbachinskiy

## Abstract

Argonaute proteins are programmable nucleases that are found in both eukaryotes and prokaryotes and provide defense against invading genetic elements. Although some prokaryotic Argonautes (pAgos) were shown to recognize RNA targets *in vitro*, the majority of studied pAgos have strict specificity toward DNA, which limits their practical use in RNA-centric applications. Here, we describe a unique KmAgo nuclease from the mesophilic bacterium *Kurthia massiliensis* that can be programmed with either DNA or RNA guides and can precisely cleave both DNA and RNA targets. KmAgo preferentially binds 16-20 nt long 5′-phosphorylated guide molecules with no strict specificity for their sequence and is active in a wide range of temperatures. In bacterial cells, KmAgo is loaded with small DNAs with no obvious sequence preferences suggesting that it can uniformly target genomic sequences. Target cleavage by KmAgo depends on the formation of secondary structure indicating that KmAgo can be used for structural probing of RNA targets. Mismatches between the guide and target sequences greatly affect the efficiency and precision of target cleavage, depending on the mismatch position and the nature of the reacting nucleic acid. These properties of KmAgo open the way for its use for highly specific nucleic acid detection and cleavage.

## Introduction

Prokaryotic Argonautes (pAgos) form a diverse family of proteins that are found in both bacteria and archaea and are related to eukaryotic Ago proteins involved in RNA interference (1–4). pAgos can be divided into three clades, including long active, long inactive pAgos and short pAgos (5). The conserved core structure of long pAgo proteins includes the N-terminal, L1, PAZ, L2, MID and PIWI domains, while short pAgos consist of only MID and PIWI domains. Long active pAgos have a catalytic tetrad DEDX (X is D, H, or K) in the PIWI domain, which binds divalent metal ions and participates in catalysis. When programmed with small nucleic acid guides, active pAgos can recognize complementary targets and perform their precise cleavage between the 10^th^ and 11^th^ positions of the guide nucleic acid (1,3,4,6).

Most previously characterized catalytically active pAgos are DNA-guided nucleases that target DNA. They were initially isolated from thermophilic bacteria, including TtAgo from *Thermus thermophilus*, PfAgo from *Pyrococcus furiosus* and MjAgo from *Methanocaldococcus jannaschii* (7–13). More recently, pAgo nucleases from mesophilic bacteria have also been described including CbAgo from *Clostridium butyricum* (14,15), LrAgo from *Limnothrix rosea* (15) and SeAgo from *Synechococcus elongatus* (16). Among them, CbAgo shows robust activity toward single-stranded DNA substrates in the range between 20 and 55 °C and can also precisely cleave double-stranded DNA at elevated temperature (14,15). Recognition of double-stranded DNA by pAgos probably requires its prior melting, which can be modulated by variations in the GC/AT-content, temperature, the action of accessory proteins or cellular processes of DNA replication, transcription and repair (8,15,17–20). In the absence of guides, some pAgos were shown to perform non-specific processing of target DNA *in vitro*, possibly resulting in autonomous generation of guide molecules, but the role of this activity *in vivo* remains to be established (13,15,16,21).

Similarly to eukaryotic Agos, pAgos have been implicated in the cell’s defense against mobile genetic elements (1,4,6). Recently, CbAgo was shown to protect bacterial cells from invaders, including plasmids and phages (17). Similarly, thermophilic TtAgo and PfAgo were shown to decrease plasmid DNA content and transformation efficiency (7,8). RsAgo from *Rhodobacter sphaeroides*, a long inactive pAgo, also preferentially targets plasmid DNA, transposons and prophages (20). In addition, TtAgo has been recently shown to participate in chromosomal DNA decatenation and completion of replication (22), suggesting that pAgos may have a wider range of cellular functions beyond defense against foreign DNA.

Ago proteins can potentially be used for specific cleavage of DNA or RNA targets, thus providing a tool for nucleic acid manipulations. However, there are only a few known examples of pAgo nucleases that can be loaded with RNA guides or recognize RNA targets. In particular, MpAgo from thermophilic *Marinitoga piezophila* binds RNA guides to cleave DNA and, with a somewhat lower efficiency, RNA targets (23). TtAgo from thermophilic *Thermus thermophilus* uses DNA guides to cleave DNA but can also process RNA (8,10,24). No mesophilic pAgos with the ability to recognize both types of nucleic acids, DNA and RNA, have been described to date. The search for other mesophilic pAgo proteins with a relaxed specificity, which could be used in both DNA and RNA fields, is therefore of a great importance. In this work, we characterize a novel mesophilic pAgo nuclease, KmAgo from *Kurthia massiliensis*, which is distantly related to CbAgo and contains the canonical catalytic tetrad in the PIWI domain (residues D527, E562, D596 and D713). We show that, similarly to other mesophilic pAgos, KmAgo is active at ambient temperatures but can be programmed with both DNA and RNA to cleave both types of nucleic acids. Furthermore, we demonstrate that KmAgo has no distinct sequence specificity toward guide or target nucleic acids both *in vitro* and *in vivo* and suggest that it can be used for site specific cleavage and structural probing of DNA and RNA.

## Results

### Expression of KmAgo in *E. coli* and analysis of associated nucleic acids

To study the biochemical properties and *in vivo* functions of KmAgo, we amplified the KmAgo gene from the genomic DNA of *K. massiliensis* and cloned it in an expression vector. We obtained *E. coli* strains with plasmid-encoded KmAgo and used them for *in vivo* assays and purification of KmAgo. The expression of KmAgo in *E. coli* led to a significant delay in cell growth in comparison with control strains grown without the addition of the inducer or containing an empty plasmid, suggesting that the protein is toxic for the cells (Fig. 1A). We purified KmAgo using metal-affinity and heparin chromatography steps, resulting in >98% pure preparations (Fig. S1, Materials and Methods) and used it for further functional analysis (see next sections).

**Fig. 1.**
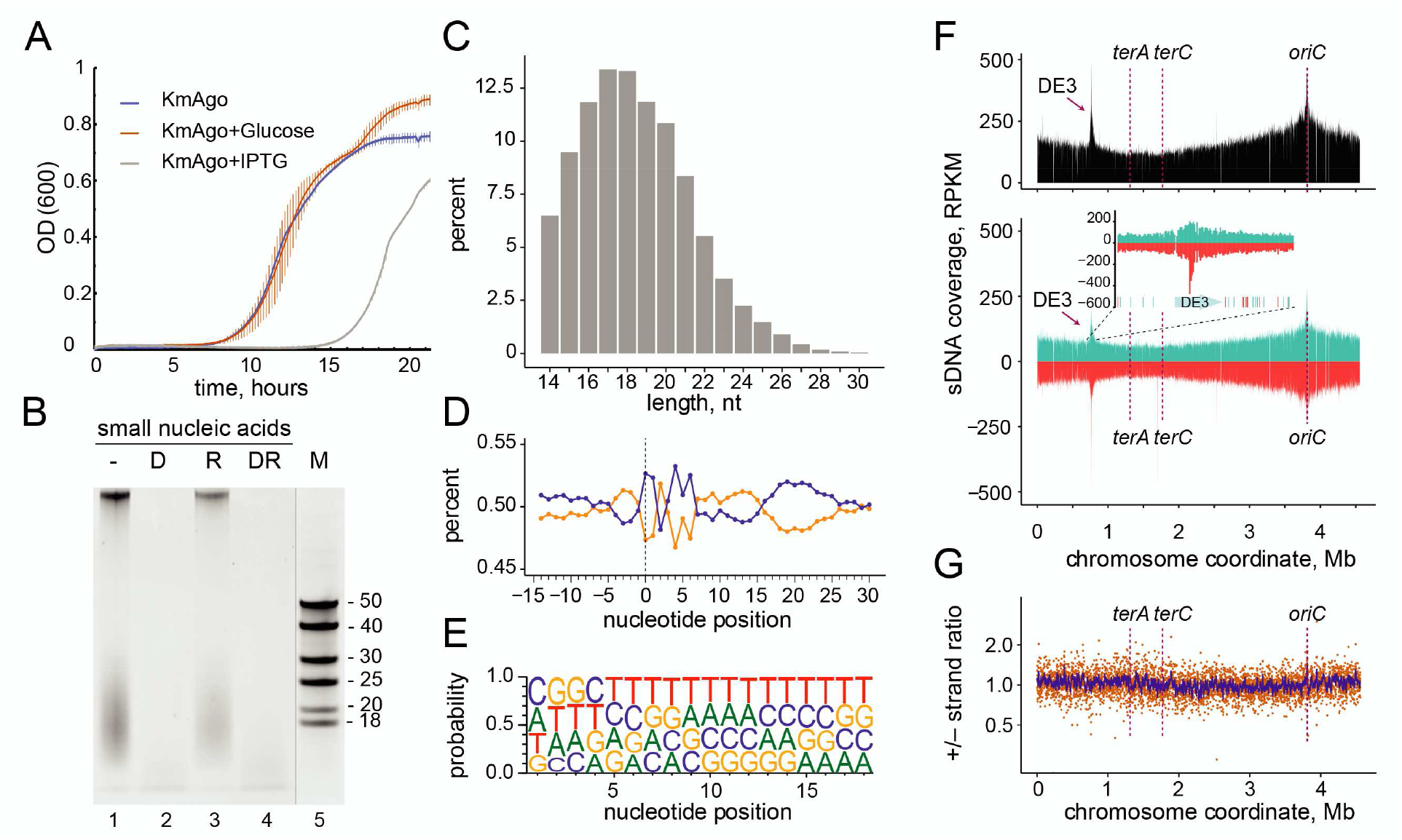
KmAgo binds small DNAs from genomic sequences during expression in *E. coli*. (A) Effect of KmAgo expression on cell growth. The experiment was performed in BL21(DE3) strains containing the expression vector (pET28-KmAgo) in the absence of IPTG, in the presence of Glc (to fully suppress possible leaky expression of KmAgo) and in the presence of the inducer (IPTG) (mean values and standard deviations from three biological replicates are shown). (B) Purification of KmAgo-associated nucleic acids after its expression in *E. coli*. Nucleic acids were treated with DNaseI (lane 2), RNase A (lane 3), both nucleases (lane 4), or left untreated (lane 1), separated by denaturing PAGE and stained with SYBR Gold. DNA length markers are shown in lane 5. (C) Length distribution of small DNAs in the sequenced libraries. (D) AT/GC-content along the smDNA sequences. Nucleotide positions relative to the smDNA 5′-end are shown below the plot (negative numbers correspond to genomic DNA adjacent to the 5′-end of smDNA). (E) Frequencies of individual nucleotides at different positions of smDNAs associated with KmAgo. (F) Genomic distribution of KmAgo-bound smDNAs mapped to the chromosome of the BL21(DE3) strain (*top*, the total number of smDNAs; *bottom*, strand-specific distribution of smDNAs; turquoise, plus strand; red, minus strand). The inset shows the genomic neighborhood of the DE3 prophage, with distribution of Chi sites in this region (turquoise, Chi sites in the plus strand; red, Chi sites in the minus strand). The numbers of smDNAs are shown in RPKM (reads per kilobase per million reads in the smDNA library). Positions of the replication origin (*ori*) and termination sites (*terA* and *terC*) are indicated. (G) Ratio of the smDNA densities for the plus and minus genomic strands. The ratio of RPKM values in each 1 kb window is shown in orange; the averaged ratio in the 10 kb window is shown in blue.

Previously studied pAgo proteins were shown to bind small guide nucleic acids when expressed in their native hosts or in the heterologous *E. coli* system (8,13,14,17,20,22). To explore if KmAgo is associated with guide nucleic acids when expressed in *E. coli* cells, we extracted and labeled nucleic acids from the KmAgo samples purified by metal-affinity chromatography. KmAgo was bound with 13-25 nt small guide DNAs (smDNAs), as could be concluded from their sensitivity to DNase I treatment and resistance to RNase A (Fig. 1B). Next, we cloned KmAgo-associated smDNAs and performed their high-throughput sequencing. As expected, the length distribution of the resulting sequences corresponded to the lengths of smDNAs isolated from KmAgo, with the peak at 16-19 nucleotides (Fig. 1C). We observed no nucleotide biases along the guide sequence, with comparable frequencies of all four nucleotides at all positions, as well as in the genomic region upstream of the guide 5′-end (Fig. 1D and E).

To analyze the origin of KmAgo-associated smDNAs, we mapped them to the *E. coli* chromosome and the expression plasmid. A significant portion of smDNAs corresponded to plasmid sequences (3.9 and 6.8 % in the two replicas). However, when normalized to the length of corresponding replicons, the ratio of smDNAs mapping to the plasmid and chromosomal DNA (24 and 44 in the two replicas) roughly equaled the known copy number of the plasmid vector (pET28, about 20 copies per cell). Thus, unlike previously studied pAgos (8,17,20), KmAgo-bound guides are not enriched in sequences derived from plasmid DNA.

Chromosomal mapping of KmAgo-bound smDNAs revealed their almost uniform distribution along the genome with no preferences for specific genomic regions, apart from a shallow enrichment around the origin of replication, which likely results from the higher DNA content in this region due to ongoing replication (Fig. 1F and Fig. S2). The smDNA reads were evenly distributed along the plus and minus genomic strands, with about equal ratio of the strands along the whole genome (Fig. 1F, G and Fig. S2). In addition to the origin region, a peak of smDNAs around the 0.77 Mb genomic coordinate was observed in one of the two replicas (Fig. 1F). The center of the peak corresponds to the DE3 prophage in the genome of the BL21(DE3) strain that was used for KmAgo expression, indicating some kinds of genomic rearrangements in this region (see Discussion). In summary, these experiments demonstrated that KmAgo is loaded with smDNA guides when expressed *in vivo*, and that these DNAs are generated by uniform sampling of chromosomal and plasmid sequences by KmAgo.

### KmAgo can use both DNA and RNA guides to target DNA and RNA *in vitro*

Most previously characterized pAgo nucleases have strict specificity toward DNA guides and DNA targets (see Introduction). To test whether KmAgo have a distinct specificity for DNA or RNA *in vitro*, we analyzed its activity using a set of synthetic guide and target oligonucleotides (Fig. 2A, Table S1). KmAgo was first loaded with guide DNA or RNA and both types of binary complexes were incubated with single-stranded DNA or RNA targets. After incubatiuon the cleavage products were analyzed by denaturing gel-electrophoresis (Fig. 2B). Surprisingly, it was found that KmAgo is active as guide-directed endonuclease with all combinations of guide and target nucleic acids, with the highest activity observed for DNA-guided DNA cleavage, and the lowest for RNA-guided RNA cleavage. Importantly, in all cases the cleavage products corresponded to the single expected cleavage site (the target position between the 10^th^ and 11^th^ nt of the guide, the canonical cleavage site observed for all previously studied Ago proteins), demonstrating that KmAgo retains high level of precision with various combinations of guides and targets. No cleavage activity was detected for a catalytically dead KmAgo variant with substitutions of two of the tetrad residues (D527A and D596A) (Fig. 2B, lane 3).

**Fig. 2.**
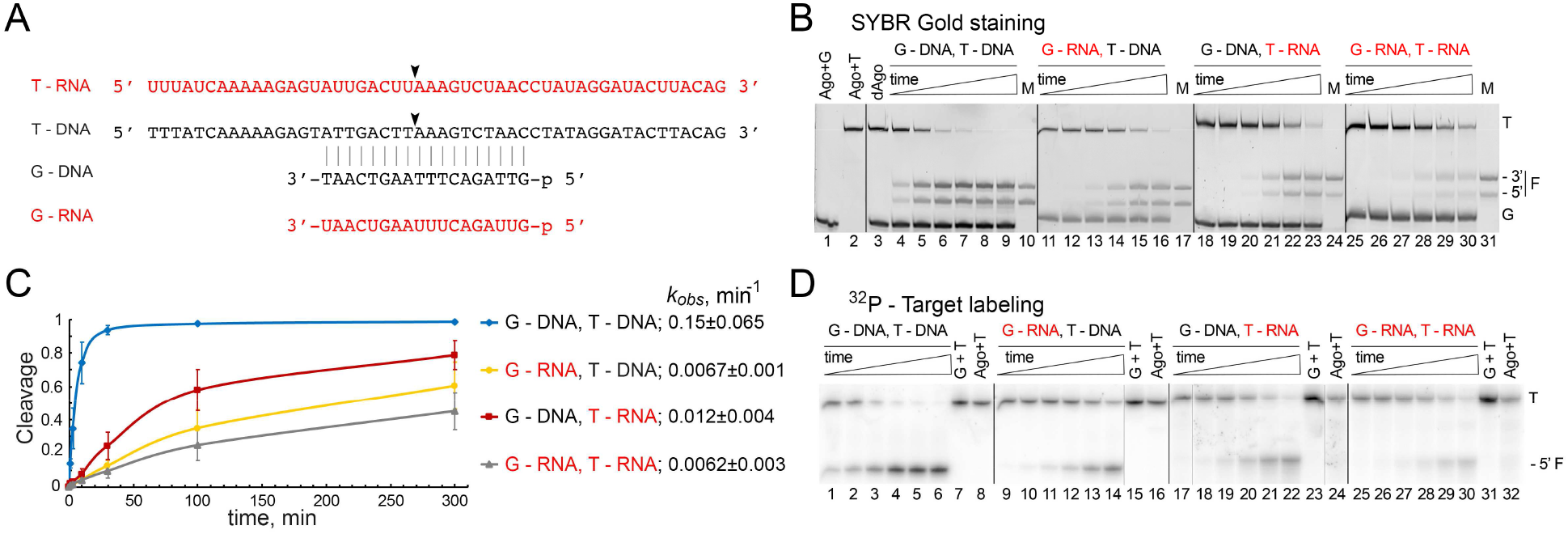
Guide and target specificity of KmAgo *in vitro*. (A) Scheme of the guide and target nucleic acids (DNA or RNA); the cleavage position is indicated with an arrow. (B) Analysis of nucleic acid cleavage by KmAgo. KmAgo pre-loaded with guide DNA or RNA was incubated with singlestranded DNA or RNA targets for indicated time intervals, nucleic acids were separated by denaturing PAGE and visualized by SYBR Gold staining. Positions of the targets (T), guides (G) and the 5′- and 3′-cleavage fragments (F) are indicated. The lengths of the synthetic RNA and DNA marker (M) oligonucleotides corresponded to the expected cleavage site between the 10^th^ and 11^th^ guide positions. (C) and (D) Kinetics of nucleic acid cleavage by KmAgo measured with radiolabeled target DNA or RNA. The data were fitted to a single-exponential equation. For each guide-target pair, the resulting *k*_obs_ value is shown (means and standard deviations from three independent measurements).

To determine the catalytic parameters of KmAgo, we measured the kinetics of the reaction using 5′-^32^P-labeled target molecules and calculated the observed rates (*k*_obs_) of target cleavage under single-turnover conditions (*i.e*. in excess of the KmAgo-guide complex over the target) (Fig. 2C and 2D). In agreement with the previous experiment, the fastest cleavage was observed for DNA guide and DNA target (*k*_obs_ = 0.15 min^−1^, corresponding to the reaction half-time t_1/2_ of about 4.6 min), followed by DNA-guided RNA cleavage (*k*_obs_ 0.012 min^−1^), and RNA-guided DNA and RNA cleavage (*k*_obs_ 0.0067 min^−1^ and 0.0062 min^−1^, respectively) (Fig. 2C). Importantly, although the reaction rates varied significantly for different guide/target combinations, at least half of the substrate was processed during the reaction time in all cases, thus making KmAgo the first Ago nuclease that is active with all variants of guide and target nucleic acids.

### KmAgo is active under a wide range of reaction conditions and has no strict sequence requirements

Divalent metal ions bound in the catalytic center of Ago proteins play the key role in the cleavage reaction. We observed that, similarly to other pAgos, the most efficient cleavage occurs with Mn^2+^ and, less efficiently, with Mg^2+^ cofactors (Fig. 3A). A low level of activity was observed with Co^2+^ and no activity was detected with Zn^2+^ and Cu^2+^. Therefore, all further experiments were performed in the presence of Mn^2+^ ions.

**Fig. 3.**
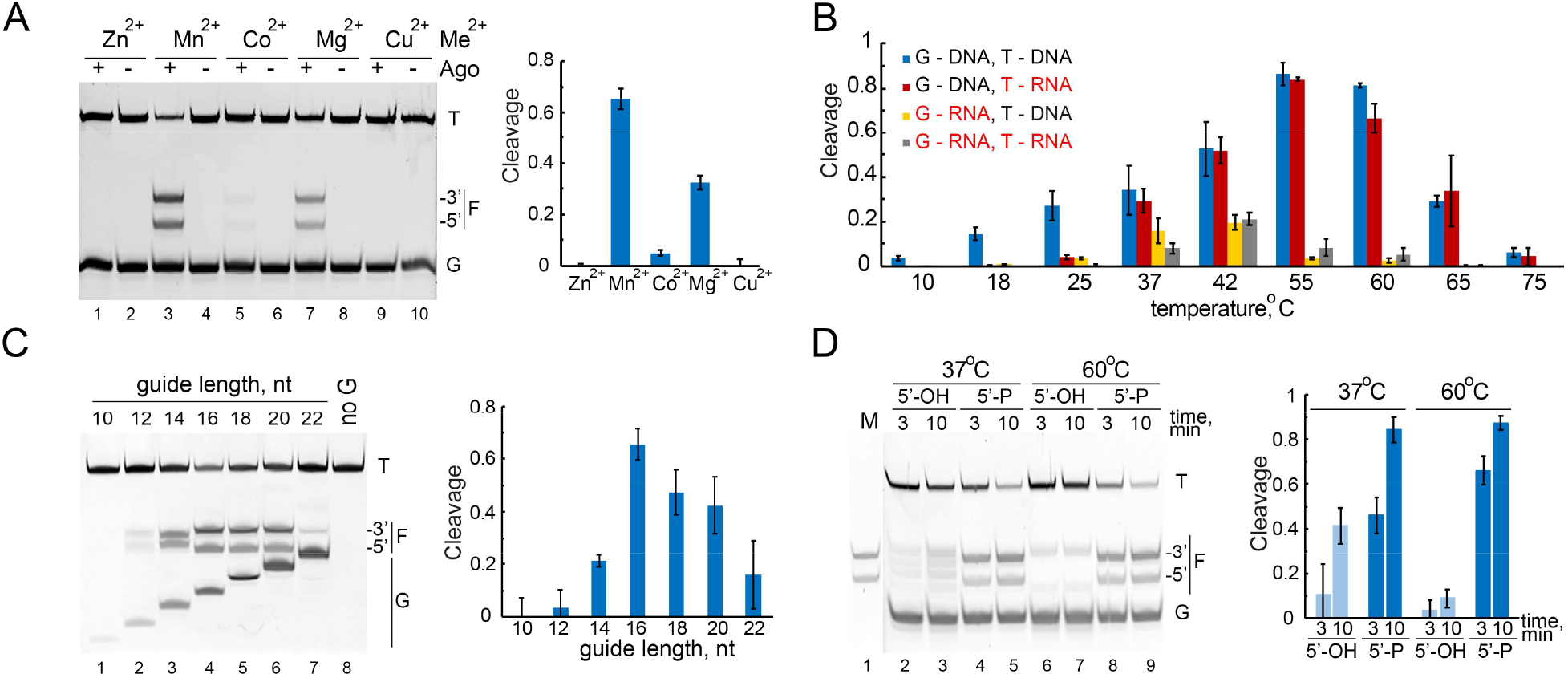
Characteristics of nuclease activity of KmAgo. (A) DNA-guided DNA cleavage by KmAgo with various divalent cation cofactors. The reaction was performed for 10 min at 37 °C. The fraction of cleaved target for each guide-target combination is shown. (B) Temperature dependence of the cleavage reaction with various combinations of guide and target nucleic acids. (C) Dependence of the efficiency of DNA cleavage on the length of guide DNA. The reaction was performed for 4 min at 37 °C. (D) The requirement of the 5′-phosphate group in guide DNA for precise target DNA cleavage. The reaction was performed for 3 or 10 min at 37 °C. The marker (M) lane contains DNA fragments corresponding to the canonical cleavage site. For each experiment, representative gels and means and standard deviations from 3-4 independent measurements are shown.

Most previously studied pAgos were isolated from thermophilic bacteria and were active only in a high temperature range (7,8,11,13,23). Recently, several DNA-guided pAgos from mesophilic bacteria were demonstrated to cleave DNA targets at ambient temperatures (14–16). We therefore determined optimal temperature conditions for nucleic acid cleavage by KmAgo. When loaded with the DNA guide, KmAgo was able to precisely cleave the corresponding DNA target at temperatures between 18 and 65 °C, with the most efficient cleavage at 37-60 °C (Fig. 3B). A similar temperature dependence was observed for the corresponding RNA target between 37 and 65 °C; however, the RNA target was cleaved less efficiently at low temperatures (18-25 °C). In the case of the RNA guide, KmAgo was active in a much narrower range of temperatures, between 37 and 42 °C, with no strong differences observed for the DNA and RNA targets (Fig. 3B). Therefore, DNA guides can be used by KmAgo to cleave both DNA and RNA targets in a wider temperature range in comparison with RNA guides.

We further studied the role of guide structure such as its length and the presence of the 5′-phosphate on target DNA cleavage. KmAgo was most active with 16-20 nt guide DNAs, with a lower cleavage efficiency observed with shorter or longer guides (Fig. 3C). Interestingly, the cleavage position was shifted by one nucleotide if shorter (12 and 14 nt) guides were used, so that the cleavage occurred between the 9^th^ and 10^th^ guide positions (Fig. 3C). Another factor that affects the precision of target processing is the presence of the 5′-phosphate in guide DNA. KmAgo can use 5′-OH guide DNA to cleave targets, however, the reaction efficiency was decreased and the cleavage positions were shifted relative to the canonical cleavage site (the cleavage occurred between 9^th^ and 10^th^ or 11^th^ and 12^th^ guide nucleotides at 37 °C, or between 11^th^ and 12^th^ nucleotides at 60 °C) (Fig. 3D).

Many studied Ago proteins from both prokaryotes and eukaryotes have certain specificity for the first nucleotide in the guide strand, which is bound in the MID pocket of Ago (20,25). In contrast, no changes in the cleavage rate and efficiency were observed when KmAgo was loaded with guides with different 5′-nucleotides but otherwise identical sequences (Fig. S3A). KmAgo was also able to cleave target sites with completely different sequences with comparable efficiency (except for one of the tested guides that was less efficient) (Fig. S3B). This correlates with the absence of strong nucleotide preferences at any guide position in smDNA associated with KmAgo *in vivo* (see above). Therefore, KmAgo can potentially be programmed for specific recognition and cleavage of any target sequence.

To test whether KmAgo is a single-turnover enzyme or a multiple-turnover enzyme that can dissociate from the target after cleavage, we analyzed the cleavage kinetics in reactions containing a 2-fold excess of target DNA over the binary guide-KmAgo complex. It was found that at 37 °C only a fraction of the target was cleaved during the 100 min course of the reaction (Fig. S4A, left panel). In contrast, when the reaction was performed at 60 °C almost all DNA was cleaved during 10 min, indicative of multiple-turnover reaction (right panel). To directly detect exchange of the reacting target molecules, we analyzed cleavage of identical DNA targets, unlabeled or labeled with a Cy3 dye, sequentially added to the reaction with the KmAgo-guide complex. The reactions were started with adding an excess of unlabeled target to the KmAgo-guide complex (2-4-fold molar ratio). After 30 minutes, Cy3-labeled target was added to the reaction. The cleavage of the labeled secondary target was detected after 3 min incubation at 60 °C, but not at 37 °C (Fig. S4B). Extended (3-hour) incubation led to processing of the labeled target at both temperatures. Thus, KmAgo acts as a multiple-turnover endonuclease, and exchange of target molecules is facilitated at high temperatures.

### Effects of guide-target mismatches on target cleavage

Precise recognition and cleavage of target nucleic acids is essential for the activities of genome defense systems that employ programmable nucleases, including CRISPR-Cas systems in prokaryotes and RNA interference systems in eukaryotes. To determine the sequence specificity of KmAgo, we analyzed the effects of mismatches between the guide and target strands on its nuclease activity, using three variants of guide and target nucleic acids: guide DNA and target DNA (Fig. 4A and Fig. S5A), guide RNA and target DNA (Fig. 4B and Fig. S5B), and guide DNA and target RNA (Fig. 4C and Fig. S5C). In the effector complexes of Ago proteins, the guide molecule can be functionally divided into several regions, including the 5′-nucleotide bound in the MID pocket of Ago, the 5′-seed region (positions 2-8), the central region surrounding the cleavage site (positions 9-12), the 3′-supplementary region (positions 13-15), and the 3′-tail region (position 16 and downstream of it) (Fig. 4). We found that mismatches in each of these regions had various effects on the efficiency and precision of target cleavage by KmAgo, depending on the nature of the reacting molecules. In reactions with DNA guides and targets (G-DNA/T-DNA), mismatches in the 3′-supplementary region of guides had the strongest effects on DNA cleavage (Fig. 4A). In addition, mismatches in the central region and at positions 4-5 in the seed region also decreased efficiency of cleavage. In contrast, when the same set of mismatched guide DNA was tested with target RNAs (G-DNA/T-RNA), no changes in cleavage were observed for mismatches in the 3′-suppplementary region (Fig. 4B). Instead, the strongest decrease in cleavage efficiency was observed for mismatches at the cleavage site and in the middle of the seed region. Finally, for reactions containing guide RNA and target DNA (G-RNA/T-DNA) the target cleavage was strongly effected by mismatches at most positions in the seed and central regions (Fig. 4C). This correlated with the lower reaction rate for this combination of guide and target nucleic acids (see above), likely making the reaction more dependent on correct base-pairing.

**Fig. 4.**
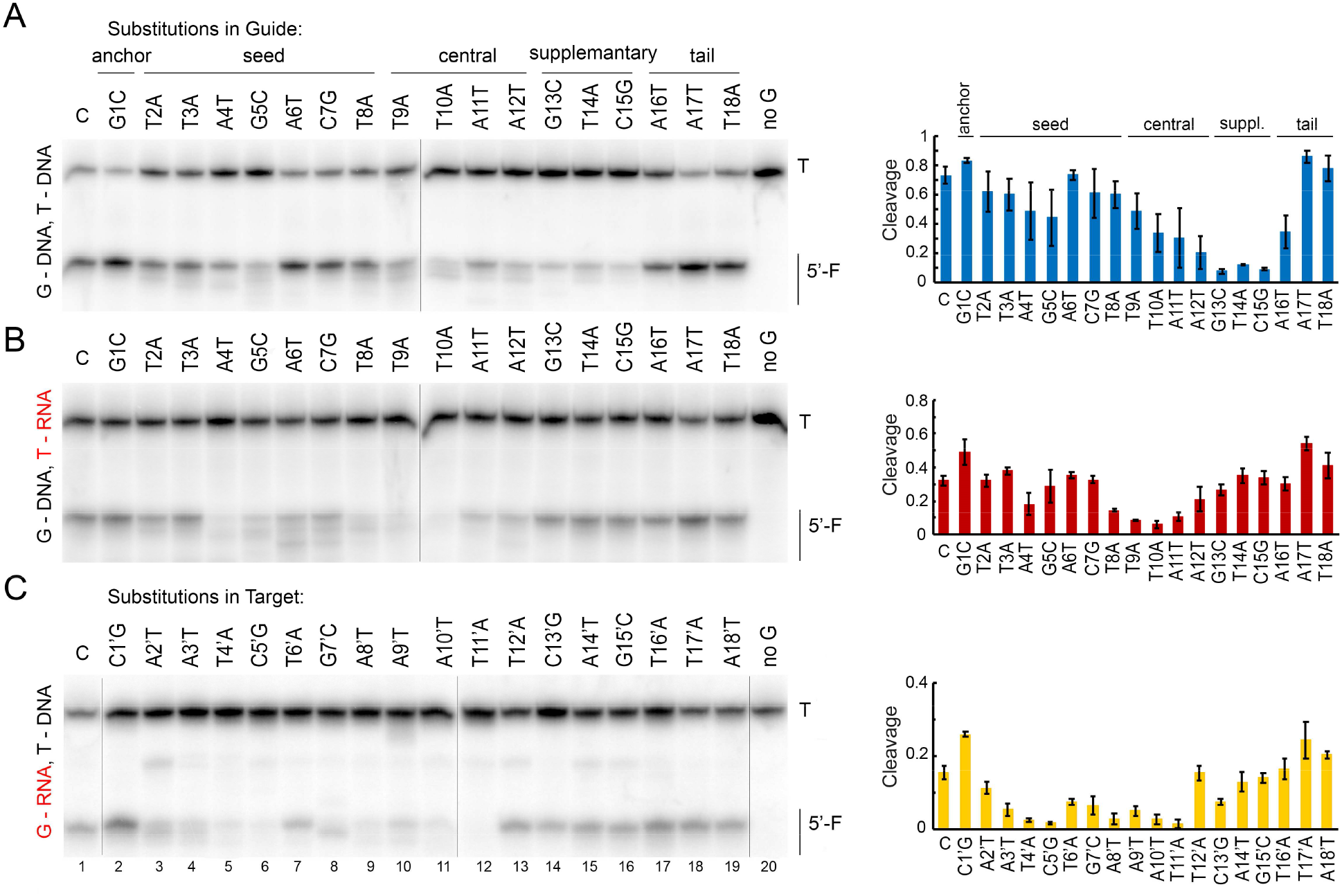
Effects of guide-target mismatches on DNA and RNA cleavage by KmAgo. Reactions with guide DNA and target DNA (A), guide DNA and target RNA (B) and guide RNA and target DNA (C) were performed at 37 °C for 15, 60 and 100 minutes, respectively. In panels A and B, a series of different DNA guides contained substitutions at each guide position, shown above panel A. In panel C, a series of DNA targets contained substitutions at each position in the region of complementarity to the guide RNA, shown above panel C (see Table S1 for full guide and target sequences). Positions of the 5′-cleavage fragments (F) are indicated. Means and standard deviations from three independent measurements.

Unexpectedly, in addition to changing the efficiency of target cleavage, mismatches also changed its pattern. The appearance of additional products resulting from cleavage at noncanonical sites was observed not only for mismatches at the site of cleavage (Fig. 4, positions 9-12) but also for mismatches in the seed region (*e.g*. at positions 2-5 in the G-DNA/T-DNA reactions, positions 2,4,5-8 in the G-DNA/T-RNA reactions, positions 2-4,7,8 in the G-RNA/T-DNA reactions) and even more surprisingly in the 3′-supplementary region (position 14 in the G-DNA/T-DNA reactions). The pattern of noncanonical cleavage fragments was highly reproducible in three independent experiments (compare Fig. 4 and Fig. S5), indicating that mismatches at individual positions have distinct effects on the conformation of the effector complex. Overall, these experiments demonstrated that the complete base-pairing between the guide and target molecules is important for precise target cleavage and that even single mismatches at certain positions can disrupt proper DNA and RNA processing by KmAgo.

### Plasmid DNA processing by KmAgo

In the experiments described above, the activity of KmAgo was tested on single-stranded DNA targets, which are preferred substrates for most previously studied pAgos (see Introduction). To determine the ability of KmAgo to process double-stranded DNA, we analyzed cleavage of supercoiled plasmid DNA with empty KmAgo or with KmAgo loaded with one or two DNA guides complementary to the two strands in the target plasmid (guides ‘g0’ and ‘g0c’, Fig. 5A). Incubation with empty or loaded KmAgo at 37 °C led to rapid relaxation of the supercoiled plasmid, resulting in disappearance of the supercoiled form and increase in the relaxed form (Fig. 5B upper panel, compare lanes 4-6 with the control sample in lane 3 in the absence of KmAgo). The presence of guides targeting the plasmid had no effect on the distribution of plasmid isoforms indicating that in these conditions both empty and loaded KmAgo can relax supercoiled DNA but cannot perform double-stranded DNA cleavage (Fig. 5B upper panel, compare lanes 7-9 and 10-12). In contrast, at 60 °C the amounts of linear plasmid DNA were higher in reactions with two guides targeting both strands of the plasmid, compared to reactions with no guides or a single guide (Fig. 5B lower panel, lanes 10-12). This result indicates that KmAgo loaded with two guides targeting both DNA strands is able to introduce double-stranded breaks into plasmid DNA at high temperatures. Some level of degradation of plasmid DNA observed with guide-free KmAgo at 60 °C (lanes 4-6 in the bottom panel) may result from the ‘chopping’ activity previously described for other pAgo proteins (13–15,21).

**Fig. 5.**
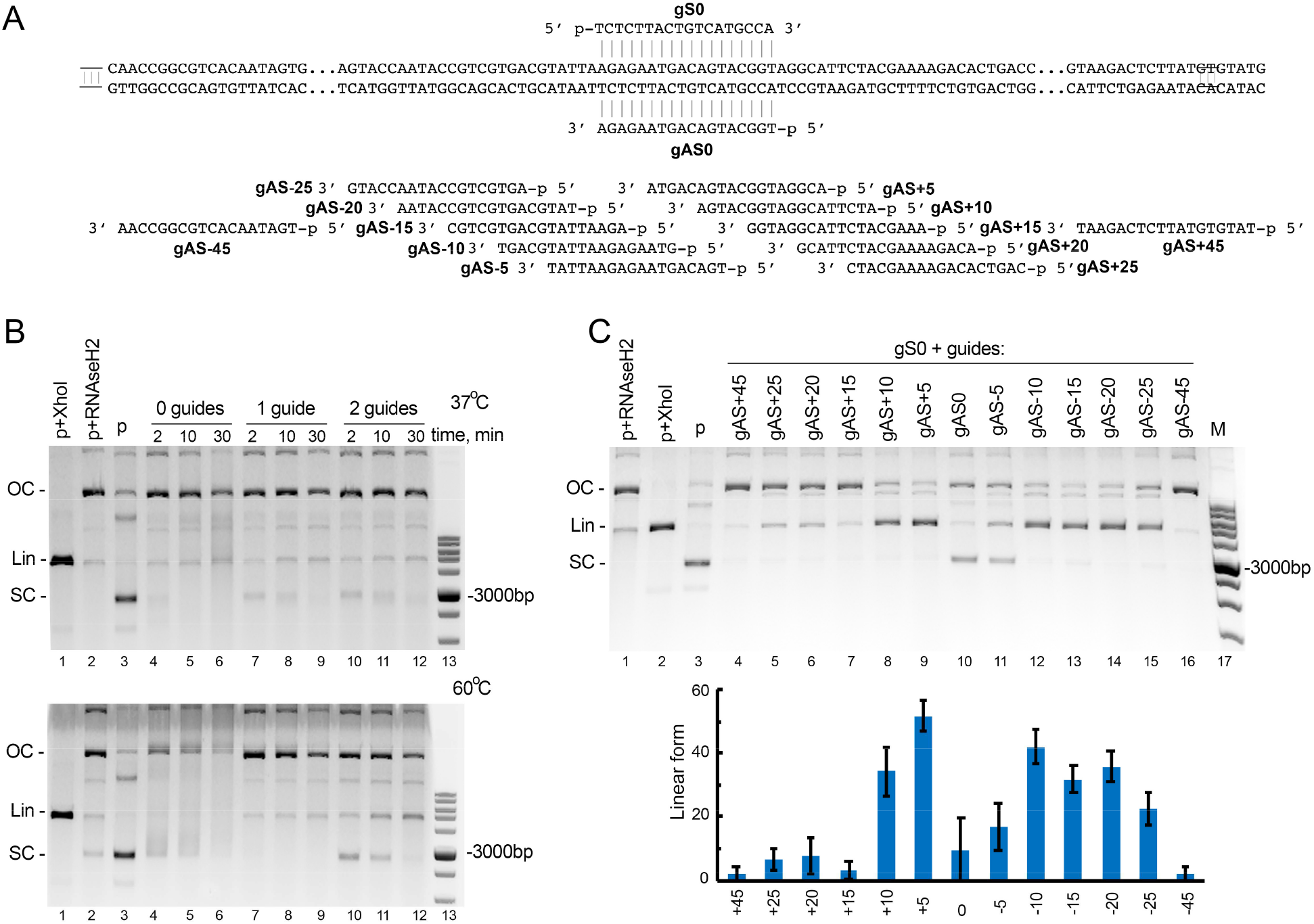
Plasmid DNA cleavage by KmAgo. (A) Scheme of the plasmid target used in the cleavage assay. Positions of guide DNAs corresponding to the sense (gS) and antisense (gAS) strands of the target are shown along the plasmid sequence. (B) The target plasmid was incubated with empty KmAgo (lanes 4-6) or KmAgo loaded with one or two guide DNAs (gS0 and gAS0; lanes 7-12) at 37 °C (top) or 60 °C (bottom) for indicated time intervals. Control samples containing linear plasmid obtained by treatment with XhoI (lane 1), relaxed plasmid obtained by treatment with RNase H2 (lane 2), and supercoiled plasmid (lane 3) were incubated in the absence of KmAgo. (C) Plasmid linearization with pairs of guide DNAs with different lengths between the cleavage sites in the two DNA strands. The reactions were performed for 2 minutes at 60 °C. Representative gels from three independent experiments are shown. Positions of supercoiled (SC), relaxed open circle (OC) and linear (Lin) plasmid forms are shown. The cleavage efficiency is shown as a fraction of linear DNA form relative to the sum of supercoiled, relaxed and linear DNA (means and averages from three measurements).

We further tested whether the relative positions of the two guide molecules targeting opposite DNA strands influence the efficiency of dsDNA cleavage. We used a set of guide pairs in which the position of the guides targeting the bottom ‘antisense’ DNA strand was shifted relative to the position of the top ‘sense’ strand guide by 5, 10, 15, 20, 25 or 45 nucleotides in both directions (g5L, g5R, g10L, g10R etc.) (Fig. 5A). Shifting the position of the ‘antisense’ guide by 5 and 10 nt to the right (Fig. 5C, lanes 8,9) or by 10, 15, 20 and 25 nt to the left (lanes 12-15) relative to the ‘sense’ guide significantly increased the efficiency of plasmid linearization in comparison with oppositely located guides (lane 10). This suggests that the effector complexes of KmAgo interfere with each other during recognition of the two DNA strands if they are located directly opposite each other.

Several previously studied pAgos were shown to decrease plasmid content and/or plasmid transformation efficiency in the host cells (including RsAgo, CbAgo, TtAgo, PfAgo) (7,8,17,20). We therefore tested whether KmAgo could have a similar effect on plasmid maintenance in *E. coli*. For this purpose, we compared the rate of plasmid loss in *E. coli* strains containing or lacking KmAgo. Under the conditions of our experiments, the expression of KmAgo did not increase the fraction of plasmid-free cells after 10 passages (Fig. S6). This is consistent with the absence of preferential loading of KmAgo with smDNAs derived from plasmid DNA (see above).

### Probing of the secondary RNA structure with KmAgo

To explore if processing of target RNA by KmAgo depends on its secondary structure, we analyzed cleavage of highly structured 6S RNA, which binds to bacterial RNA polymerase and serves as a regulator of gene expression at the stationary phase (Fig. 6A) (26). KmAgo was loaded with guide DNAs corresponding to different regions of 6S RNA that either maintain single-stranded conformation or are involved in extensive base-pairing (6S-1 through 6S-10; Fig. 6A). The cleavage products were observed with all guide DNAs when reactions were performed at 60 °C (Fig. 6B). In contrast, at 37 °C the cleavage efficiency greatly varied for different target positions. The most efficient cleavage was observed with guides 6S-2 and 6S-3 corresponding to the central singlestranded bubble in 6S RNA (Fig. 6C, lanes 2 and 3). In contrast, no cleavage could be detected with guides 6S-1 and 6S-6, corresponding to fully double-stranded target regions. The efficiency of cleavage was also significantly lower for other guides corresponding to partially double-stranded target regions. For some of these guides (6S-4, 6S-5, 6S-9) the RNA region corresponding to the seed segment of guide DNA was directly involved in base-pairing with other segments of 6S RNA, while for others (6S-7, 6S-8) the seed region was partially exposed but the target site could not be cleaved efficiently. Taken together, these experiments indicate that base-pairing of a target RNA site determines the efficiency of its processing by KmAgo, while destabilization of the secondary structure at elevated temperatures allows for efficient processing at all sites. Thus, KmAgo can potentially be used for precise processing and structural probing of RNA targets.

**Fig. 6.**
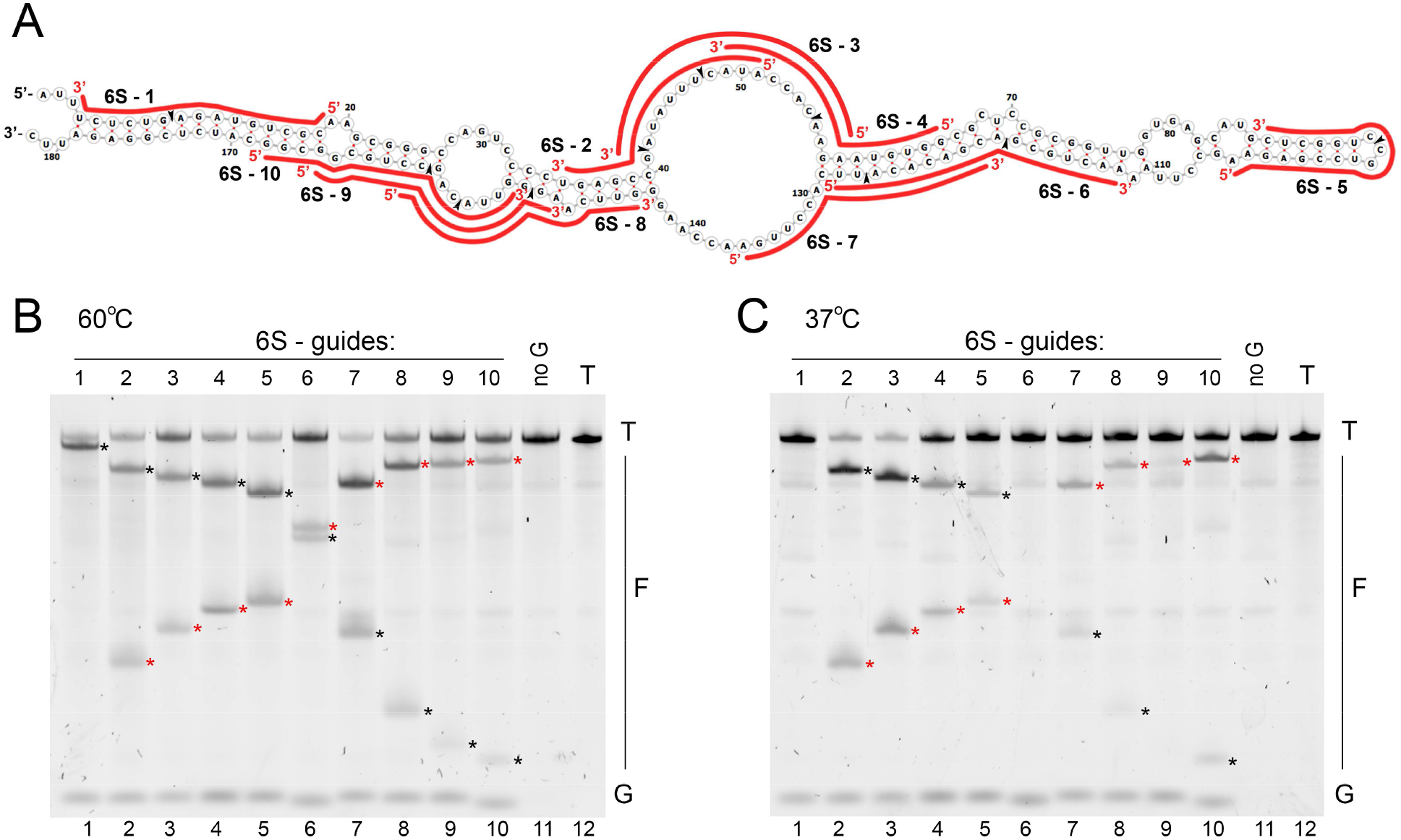
RNA probing with KmAgo. (A) Secondary structure of 6S RNA used in the experiments. Positions of small guide DNA loaded into KmAgo are shown with red lines (see Table S1 for full guide sequences). Positions of the expected cleavage sites for each guide smDNA are indicated with arrowheads. (B) Analysis of the cleavage products obtained after incubation of KmAgo with 6S RNA for 30 min at 60 °C. (C) The same experiment performed at 37 °C. Representative gels from two independent experiments are shown. Positions of the 5′-terminal and 3′-terminal cleavage fragments are indicated with red and black asterisks, respectively.

## Discussion

In this study, we have characterized a novel pAgo nuclease from the mesophilic bacterium *K. massiliensis*. In comparison with known pAgos KmAgo has an unusually broad specificity toward guide and target nucleic acid molecules and can target both DNA and RNA at physiological temperatures. It is therefore the first example of an Ago protein which is able to utilize all combinations of nucleic acids. Furthermore, we have shown that KmAgo can target singlestranded DNA and RNA under a wide range of reaction conditions, with the efficiency and precision of cleavage being modulated by temperature, divalent ion cofactors, the length and the structure of the small nucleic acid guide and its complementarity to the target.

In bacterial cells, KmAgo is associated with small guide DNAs of varying sequences suggesting that DNA is its natural substrate *in vivo*. Analysis of KmAgo-bound smDNAs have revealed no obvious preferences for specific genomic regions, except for some enrichment around the replication origin, likely as a result of a higher DNA content in this region in the dividing bacterial population. Interestingly, in one of two replicate experiments we have also observed a peak in smDNAs corresponding to the DE3 prophage in the BL21(DE3) chromosome. A similar peak of smDNAs around DE3 was observed previously during analysis of smDNAs associated with CbAgo in *E. coli* in the stationary phase of growth (17). This suggests that this chromosomal DNA region may be amplified in *E. coli* cells expressing KmAgo, possibly as a result of partial prophage excision during the stationary stage, which may promote local DNA repair and replication.

At the same time, we have not observed peaks of KmAgo-associated smDNAs in other genomic regions, particularly at the sites of replication termination, which are specifically targeted by CbAgo and TtAgo (17,22). Furthermore, KmAgo has no preference for smDNAs derived from the expression plasmid and does not affect plasmid maintenance in *E. coli*. This contrasts previous results obtained with other pAgos, including TtAgo, PfAgo, CbAgo and MjAgo which were shown to preferentially target plasmid DNA and decrease plasmid content and transformation efficiency (7,8,13,17). CbAgo can also induce DNA interference between homologous plasmid and chromosomal regions resulting in targeted chromosomal DNA processing (17). In contrast, KmAgo appears to passively capture smDNAs possibly processed from cellular DNA by other nucleases resulting in their uniform distribution along the chromosomal and plasmid sequences. It remains to be established whether KmAgo can directly participate in DNA processing or be programmed for specific DNA recognition *in vivo*.

*In vitro*, KmAgo preferentially uses 16-20 nt long 5′-phosphorylated guide DNAs and is mostly active with manganese or magnesium ions, similarly to previously studied pAgos. However, in contrast to several other pAgos, including TtAgo, MjAgo and RsAgo (8,11,20,27), KmAgo has no strong sequence preferences toward the 5′-guide nucleotide or any other position within the guide sequence. This is consistent with the broad spectrum of smDNAs associated with KmAgo in bacterial cells and potentially allows specific targeting of any desired sequence. Importantly, even single-nucleotide mismatches with the target molecule greatly affect the nuclease activity of KmAgo. We have shown that the efficiency and the position of cleavage can be modulated by mismatches in the seed, central and 3′-supplementary regions of the guide, and that cleavage of DNA or RNA targets is mostly affected by mismatches at different positions. The presence of mismatches also changes the precision of target cleavage by KmAgo. Surprisingly, these effects have been observed not only for mismatches immediately at the site of cleavage but also for many positions inside the seed or 3′-supplementary region, suggesting that these mismatches strongly affect the conformation of the catalytic complex and dislodge the target molecule from its correct position in the active site. In comparison, mismatches between the guide and target nucleic acids did not strongly affect the cleavage site for TtAgo or the structure of the ternary complex of RsAgo (9,27).

Given the strong effects of mismatches on KmAgo-dependent cleavage, accurate design of guide oligonucleotides may allow highly specific discrimination of even closely related target sequences by KmAgo in various applications. In particular, specific degradation of one of the target variants can be used for enrichment of rare nucleic acid variants, both DNA and RNA, for their further detection. Thermophilic TtAgo and PfAgo were previously applied for precise recognition and cleavage of complementary targets and their closely related variants depending on the presence of mismatches (28,29). Similarly, thermophilic MpAgo, programmed with RNA guides, was used for selective binding of rare RNA variants with single-nucleotide changes without their cleavage (30). In comparison with thermophilic proteins, KmAgo offers a more flexible tool for the detection of single-nucleotide variations in the target nucleic acids because it is active in a wider range of temperatures. The specificity of KmAgo can be likely further increased by using RNA guides that are more sensitive to the presence of mismatched nucleotides.

All analyzed pAgo proteins, including KmAgo, are significantly less active on dsDNA than on ssDNA substrates. Similarly to KmAgo, it was shown previously that TtAgo, CbAgo and RsAgo can interact with double-stranded DNA only if its melting is facilitated by various factors, including low GC-content, increased temperature, or supercoiling (8,14,15,18,21). However, since KmAgo retains high level of activity up to 60 °C, it can be used for site-specific DNA cleavage under these conditions.

Finally, we have demonstrated that KmAgo can be used for specific cleavage and structural probing of RNA targets. We have found that the efficiency of cleavage depends on the presence of double-stranded regions in the RNA target and that the secondary structure formation prevents RNA cleavage by KmAgo. This is consistent with the inability of KmAgo to efficiently cut doublestranded DNA at ambient temperatures. Thus, KmAgo-dependent RNA cleavage can allow detection of the conformational state of RNA targets. Previously, eukaryotic Ago protein from the budding yeast *Kluyveromyces polysporus* was used for site-specific RNA cleavage (31). Similarly to our findings, it was found that the efficiency of cleavage depended on the secondary RNA structure. Importantly, in comparison with previous works we have shown that KmAgo can also perform efficient RNA cleavage at elevated temperatures that promote melting of the secondary structure and thus make possible uniform RNA modification. Therefore, KmAgo can also be used for precise cleavage of complex RNA targets independently of the secondary structure formation. Furthermore, KmAgo can potentially be applied in other RNA-centric methods such as recently developed Ago-based fluorescence *in situ* hybridization (Ago-FISH) (32). In conclusion, we demonstrate that KmAgo is a unique programmable nuclease that has a broad specificity to guide and target nucleic acids and can potentially be used in a wide range of applications in the field of nucleic acid biotechnology.

## Methods

### Protein expression and purification

The gene of KmAgo (WP_010289662.1) was amplified by PCR from genomic DNA of *Kurthia massiliensis* JC30 and cloned into the pET28b expression vector in frame with the N-terminal His_6_-tag using the NheI-XhoI restriction sites. *E. coli* BL21(DE3) was transformed with the expression plasmid and the cells were cultivated in the LB medium with 50 μg/ml kanamycin at 25 °C until OD_600_=0.4, cooled down to 16 °C, induced with 0.2 mM IPTG and grown at 16 °C overnight with aeration. The cells were collected by centrifugation and stored at −70 °C. The cell pellet was resuspended in buffer A (30 mM Tris-HCl pH 7.9, 0.3 M NaCl, 5% glycerol) supplemented with 1 mM of PMSF and disrupted using Cell Disruptor CF (Constant Systems). The lysate was cleared by centrifugation, and loaded onto Co^2+^-charged TALON Metal Affinity Resin (Clontech) for 1.5 hours with agitation. The beads were washed with buffer A containing 5 mM of imidazole, than with the same buffer with 20 mM of imidazole, and eluted with buffer containing 300 mM imidazole. The eluted protein was diluted 3-fold with a buffer containing 20 mM Tris-HCl pH 7.9 and 5% glycerol. The protein sample was loaded onto a Heparin FF column (GE Healthcare) equilibrated with 20 mM Tris-HCl pH 7.9, 100 mM NaCl and 5% glycerol, washed with 10 volumes of the same buffer and eluted with a linear NaCl gradient (0.1 – 1.0 M). The purity of the final protein samples was assessed by denaturing PAGE with Coomassie staining. Fractions containing KmAgo were concentrated using Amicon Ultra 50K (Merck Millipore), placed in a storage buffer (40 mM Tris– HCl, 0.3 M NaCl, 50% glycerol, 1mM DTT, 0.5 mM EDTA, pH 7.9), aliquoted and frozen in liquid nitrogen. The protein concentration was determined by the Qubit protein assay kit (Thermo Fischer Scientific). Catalytically dead mutant variant (D527A; D596A) was obtained by site-directed mutagenesis using QuikChange Lightning Multimutagenesis kit (Agilent), expressed and purified in the same way.

### Analysis of KmAgo-associated smDNAs

Small nucleic acids were extracted by phenol-chloroform treatment from KmAgo after the first purification step (Co^2+^-resin). Nucleic acids were treated with DNase I or RNase A and analyzed by PAGE as described previously (15). Small DNA libraries were prepared as described in (17). Briefly, extracted nucleic acids were ethanol-precipitated, treated with RNase A (ThermoFisher), phosphorylated with polynucleotide kinase (New England Biolabs), ligated with adaptor oligonucleotides and sequenced using the HiSeq2500 platform (Illumina) in the rapid run mode (50-nucleotide single-end reads). Analysis of smDNA sequences was performed as described previously (17).

### Analysis of cell growth

Overnight culture prepared from *E. coli* BL21(DE3) transformed with the expression plasmid pET28-KmAgo or the empty vector was diluted 200 times until OD 0.015 in the LB medium containing 50 μg/ml kanamycin and 0.2 mM IPTG or 0.2% glucose and aliquoted in a 24-well plate 1 ml per well. The plate was incubated at 30 °C with agitation at 200 rpm and OD was measured each 10 minutes in a Tecan microplate reader. Data from three independent biological replicates were used to build the growth curves.

### Single strand nucleic acid cleavage

The cleavage assays were performed using synthetic guide and target DNAs and RNAs (see Table S1 for oligonucleotide sequences). For some experiments 5′-Cy3- or 5′-^32^P-labeled targets were used. The 6S RNA gene was PCR-amplified from the genomic DNA of *E. coli* strain MG1655 including the T7 RNA polymerase promoter in the forward primer. 6S RNA was synthesized by T7 RNA polymerase (Thermo Fisher Scientific), purified by 8% denaturing PAGE, treated with DNase I and ethanol precipitated. The cleavage reactions were performed in low adhesion tubes at 37 °C in a buffer containing 20 mM Tris-HCl pH 7.4, 100 mM NaCl, 10% glycerol, and 10 mM MnCl_2_. To analyze the effect of various divalent cations 10 mM MgCl_2_, CoCl_2_, CuCl_2_ or ZnCl_2_ were added instead of MnCl_2_. 500 nM of KmAgo was mixed with 500 nM guide DNA or RNA and incubated for 10 min at 37 °C for guide loading. All guides were 5′-phosphorylated using T4 PNK (New England Biolabs) except experiments with 5′-OH guides. Target DNA or RNA was then added to the final concentration of 100 nM. For analysis of temperature dependence of ssDNA cleavage, KmAgo was loaded with guide DNA for 10 min at 37 °C, the samples were transferred to indicated temperatures in a dry block heater (BioSan, CH-100), target DNA or RNA was added and the samples were incubated for 4 minutes with G-DNA and T-DNA, 40 minutes with G-DNA and T-RNA and 60 minutes for G-RNA with T-DNA or T-RNA. For experiments on multiple turnover cleavage (Fig. S4A) an excess of target DNA (200 nM) was incubated with 100 nM KmAgo-guide complex (equimolar KmAgo and guide ratio) at 37 °C or 60 °C. For experiments in Fig. S4B, 50 or 100 nM of the KmAgo-guide complex was incubated for 30 min with 200 nM of label-free target DNA and then 5′-Cy3 labeled target DNA was added to 200 nM concentration. For analysis of 6S RNA cleavage, the reactions were performed for 30 minutes with corresponding DNA guides. All reactions were carried out at 37°C or 60°C as indicated. The reactions were stopped after indicated time intervals by mixing the samples with equal volumes of stop solution (8 M urea, 20 mM EDTA, 0.005% Bromophenol Blue, 0.005% Xylene Cyanol). The cleavage products were resolved by 19% denaturing PAGE, stained with SYBR Gold (Invitrogen), visualized with a Typhoon FLA 9500 scanner (GE Healthcare), and analyzed by the ImageQuant (GE Healthcare) software. For reactions containing Cy3-labels, the gels with the reaction products were first scanned in the Cy3 channel and then stained with SYBR Gold. In the case of 6S RNA, the cleavage products were resolved by 10% denaturing PAGE and stained with SYBR Gold.

### Plasmid DNA cleavage

For analysis of plasmid DNA cleavage, the final pAgo and guide concentrations were 500 nM. When using two guide molecules, two samples of KmAgo were independently loaded with the guides (at the 1:1 KmAgo:guide ratio) and then mixed together to the final concentration of 500 nM. The sequences of all guide oligonucleotides are shown in TableS1. The target plasmid used in the assays is a pJET1.2 derivative (Thermo Fisher Scientific). The plasmid was added to the reaction mixtures to the final concentration of 2 nM, followed by incubation for indicated time intervals at 37 or 60 °C. The reactions were stopped by treatment with Proteinase K for 20 minutes at 25 °C, the samples were mixed with 6× SDS-free Purple Loading Dye (New England Biolabs) supplemented with SYBR Gold and the cleavage products were resolved by native 1.2% agarose gel electrophoresis. The relaxed plasmid in control samples was generated by treatment of the supercoiled plasmid with RNase H2 (New England Biolabs) for 2 hours at 37°C, the linear plasmid was obtained by treatment with a single cut restriction endonuclease (XhoI, Thermo Fisher Scientific).

## Acknowledgements

We thank Sergei Ryazansky for help with bioinformatic analysis of pAgo proteins and Denis Yudin for initial analysis of smDNA libraries. This study was in part supported by the Russian Foundation for Basic Research (18-29-07086) and Russian Science Foundation (grant 19-14-00359).

## Supplementary Information

**Fig. S1.**
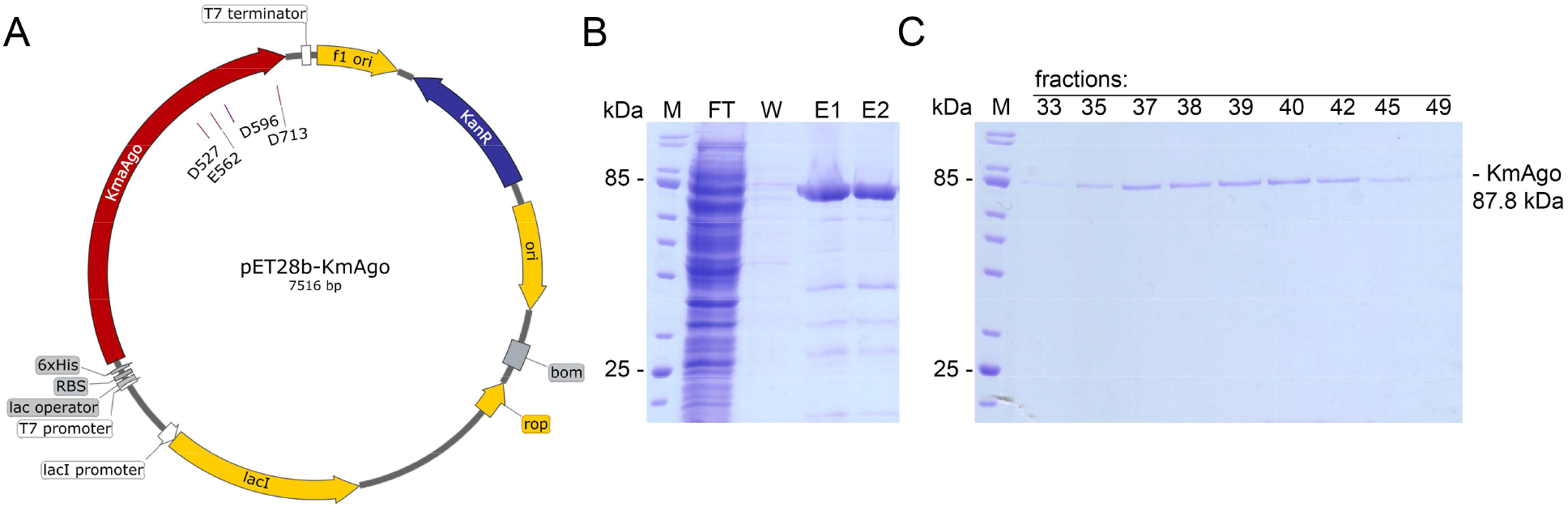
Purification of KmAgo from *E. coli* cells. (A) Scheme of the expression plasmid. Positions of the catalytic tetrad residues in KmAgo are shown. (B) The first purification step by Co^2+^-affinity chromatography. M, molecular weight marker; FT, flowthrough fraction; W, washing step with buffer containing 20 mM imidazole; E1 and E2 – elution fractions. (C) The second purification step by Heparin affinity chromatography (Coumassie staining). Fraction numbers after elution with a 0.1-1.0 M NaCl gradient. The position of KmAgo is indicated.

**Fig. S2.**
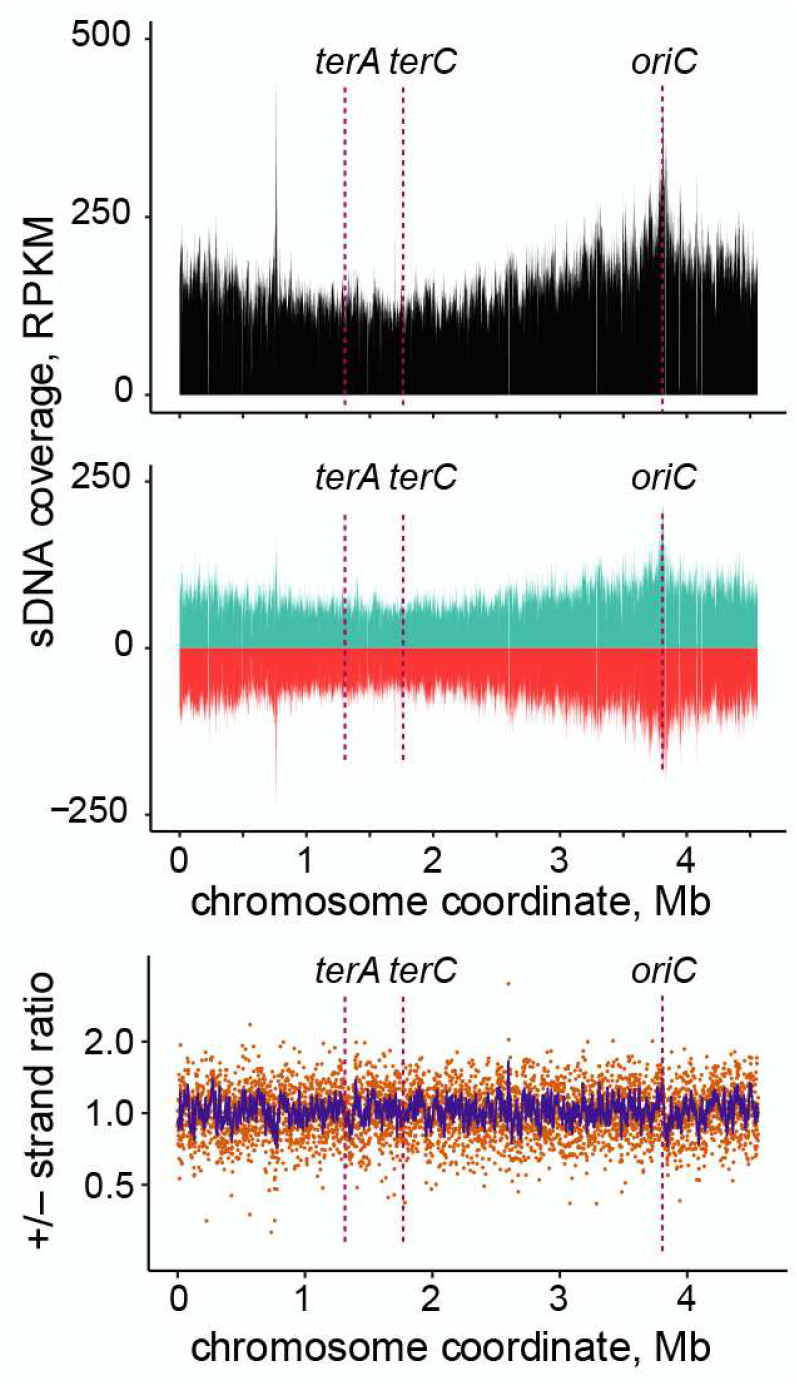
Second replica of KmAgo-associated smDNAs mapping to the chromosome of *E. coli* BL21(DE3). Positions of the origin of replication (*oriC*) and replication termination sites (terA and terC) are indicated along the chromosomal coordinate. SmDNA coverage is shown in RPKM.

**Fig. S3.**
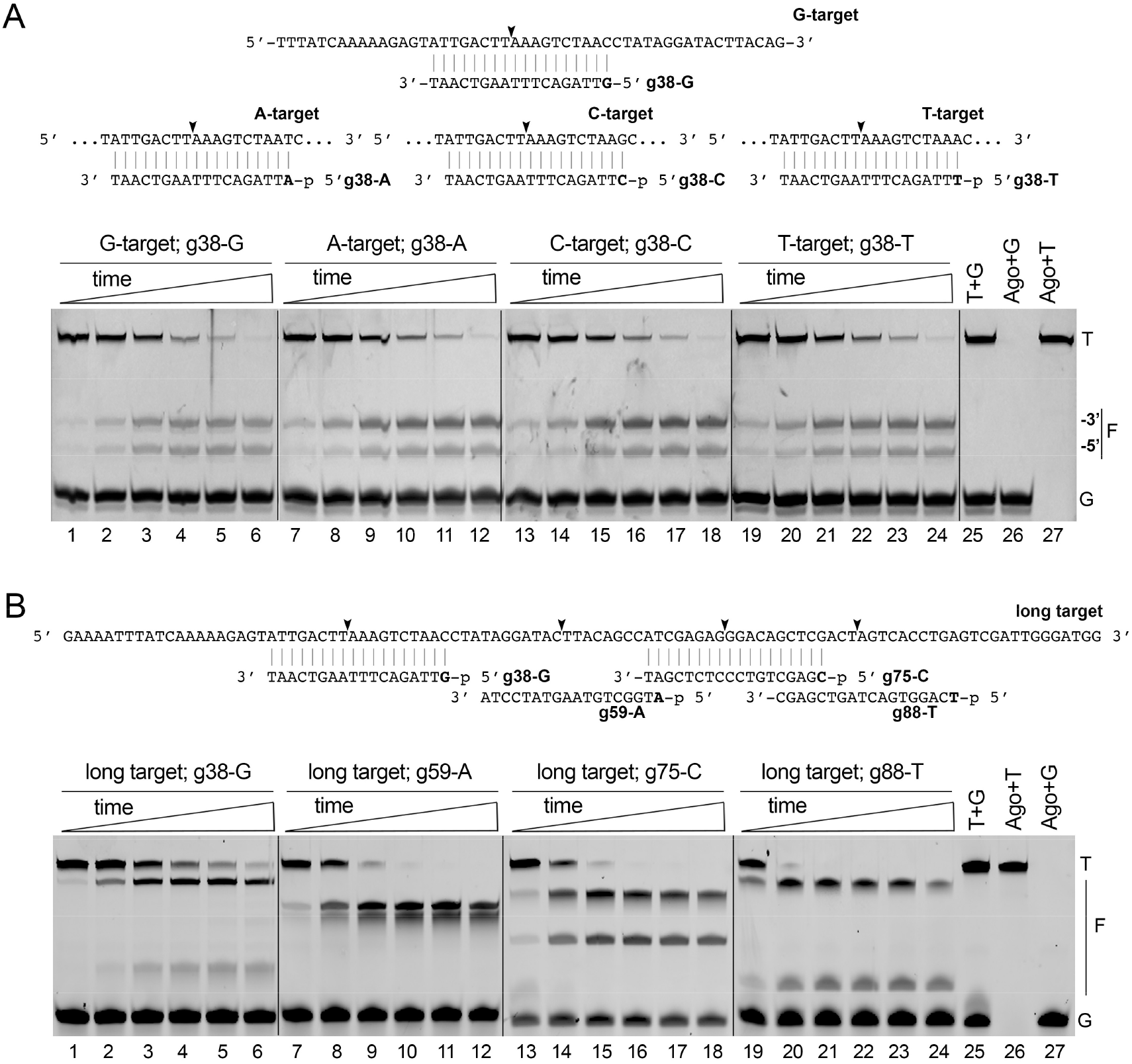
Analysis of single-stranded DNA cleavage by KmAgo loaded with various guide DNA sequences. (A) Kinetics of cleavage of the 50 nt target by KmAgo loaded with guide DNAs containing different 5′-nucleotides (G, A, C, and T) and corresponding to the same target site. The scheme of the target and guide DNAs is shown on the top. (B) Kinetics of cleavage of a longer target variant (shown on the top) with guide DNAs containing different 5′-nucleotides and corresponding to different target sites. Note that the rate of the reaction with the ‘-38’ guide DNA is somewhat lower than in the case of other guide molecules. Positions of the target (T), guide (G) molecules and the cleavage fragments (F) are indicated.

**Fig. S4.**
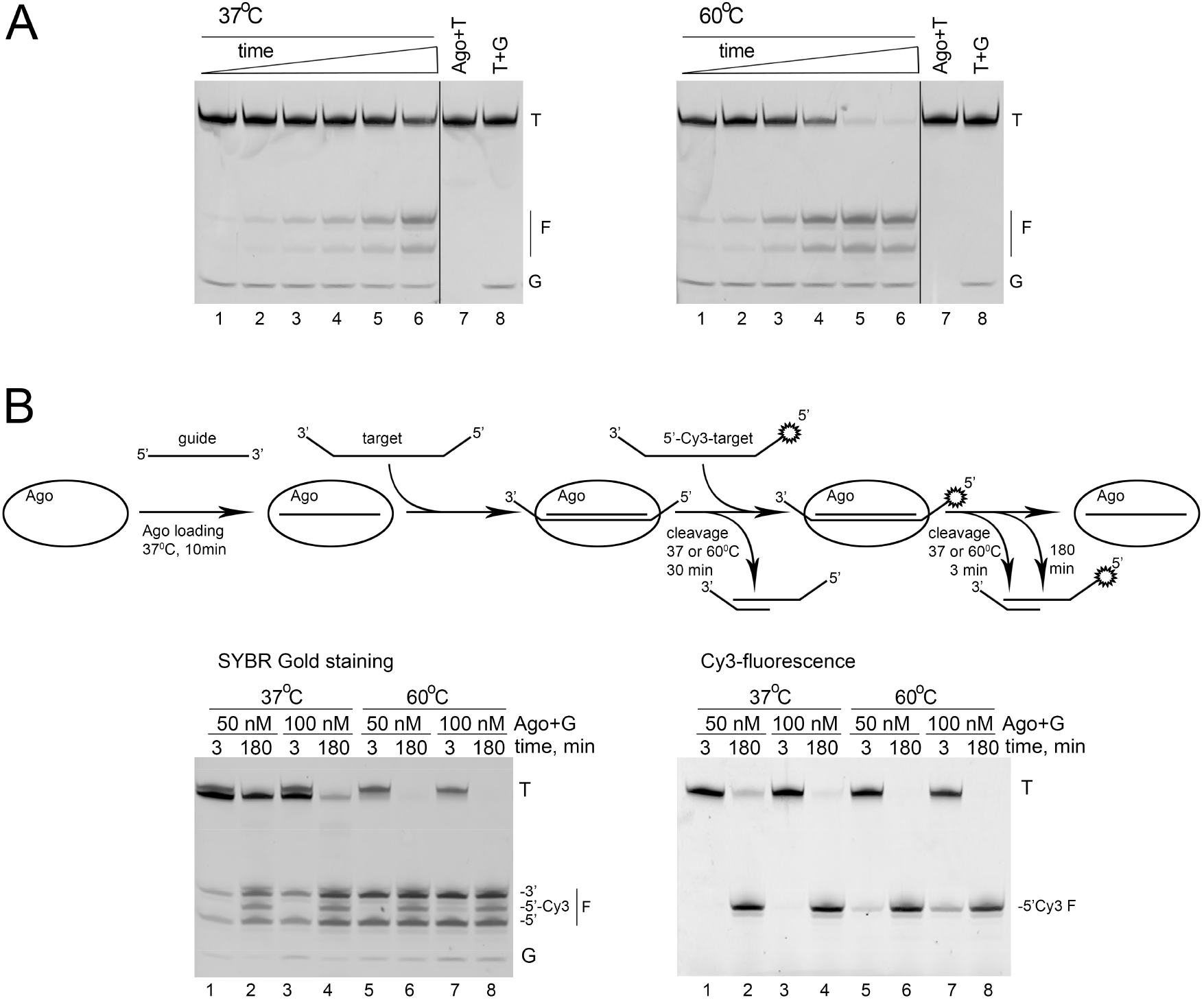
Multiple turnover DNA cleavage by KmAgo. (A) Comparison of the kinetics of target DNA cleavage by KmAgo at different temperatures. KmAgo (100 nM) pre-loaded with guide DNA (100 nM) was incubated with target DNA (200 nM) at 37 °C (left panel) or 60 °C (right panel) for indicated time intervals. (B) Experiments on two-step target DNA cleavage. KmAgo (50 nM or 100 nM) pre-loaded with equimolar amounts of guide DNA was incubated with unlabeled target DNA (200 nM) for 30 min at 37 °C, followed by the addition of identical 3′-Cy3-labeled target DNA (200 nM). The reactions were stopped after 3 minutes or 3 hours, separated by denaturing PAGE and visualized by SYBR Gold staining (left) or Cy3 fluorescence scanning (right). Positions of the target (T), guide (G) molecules and cleavage fragments (F) are indicated. After 3 minutes at 37 °C, only the first unlabeled DNA target is cleaved (lanes 1,3, with no Cy3-labeled products visible on lanes 9,11). At 60 °C initial cleavage of the Cy3-labeled target is also visible (lanes 13,15). After 3 hours, the Cy3-containing target is cleaved with high efficiency at both temperatures (lanes 10,12,14 and 16).

**Fig. S5.**
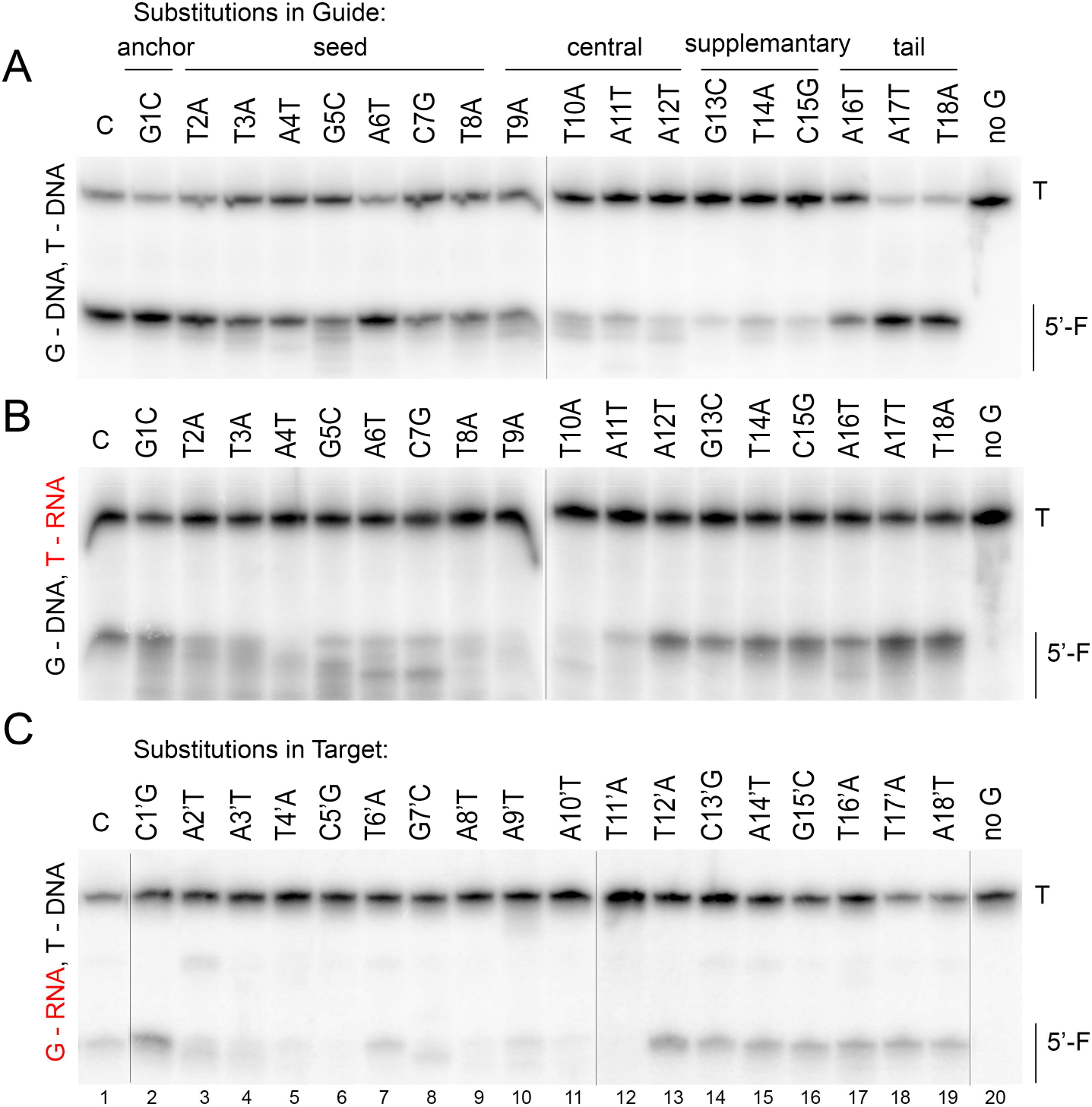
Reproducibility of the target DNA or RNA cleavage by KmAgo with mismatched guide molecules. A replica of the experiment from Fig. 4 is shown.

**Fig. S6.**
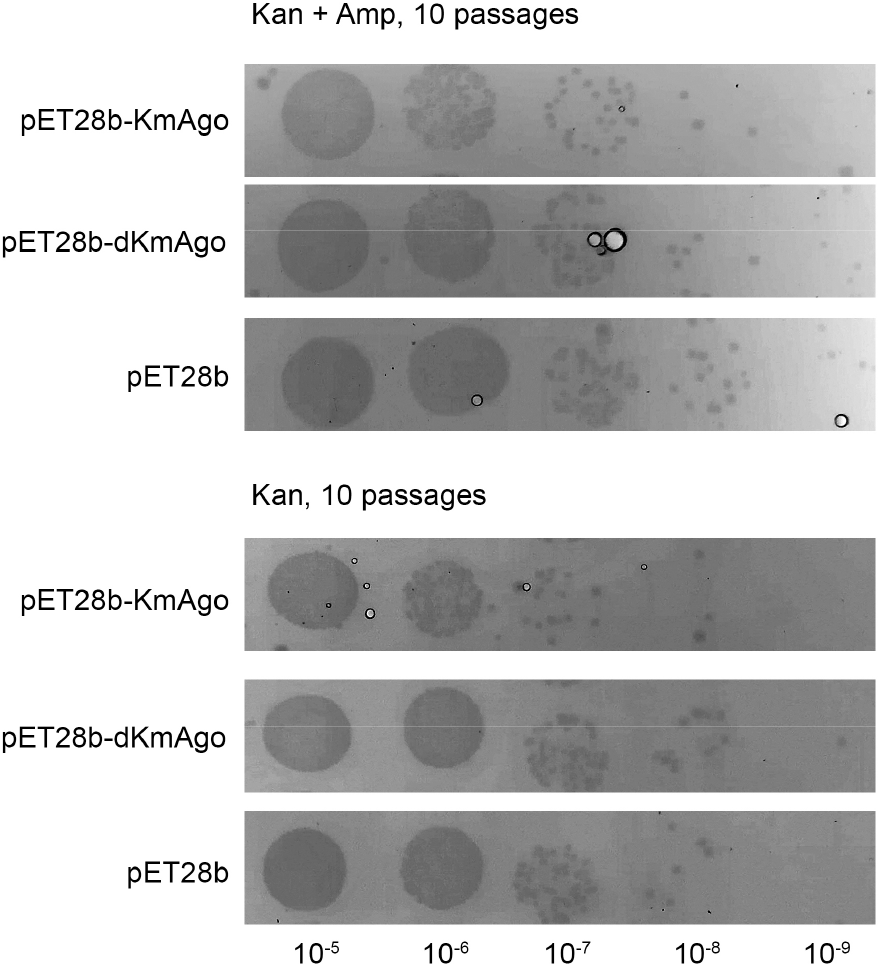
Effects of KmAgo on plasmid loss. *E. coli* BL21(DE3) were co-transformed with pSRKAmp (Amp^R^) and pET28b-KmAgo or pET28b-dKmAgo (catalytically dead Ago) or empty vector pET28b (Kan^R^). The cells were cultivated in the LB medium with 50 μg/ml kanamycin and 0.2 mM IPTG in the absence of ampicillin (to allow the loss of the pSRKAmp plasmid) or with kanamycin, IPTG and 200 μg/ml ampicillin (for control) at 18°C during 10 passages. The duration of each passage was 24h; after 24 h the cells were diluted 1:100 in the fresh LB medium with IPTG and antibiotics. After the final passage, serial dilutions of bacterial cultures were plated on agar LB dishes with kanamycin and ampicillin to compare the number of cells containing both plasmids and incubated at 37 °C overnight. No differences could be seen between the cultures that were grown with and without ampicillin.

**Table S1.**
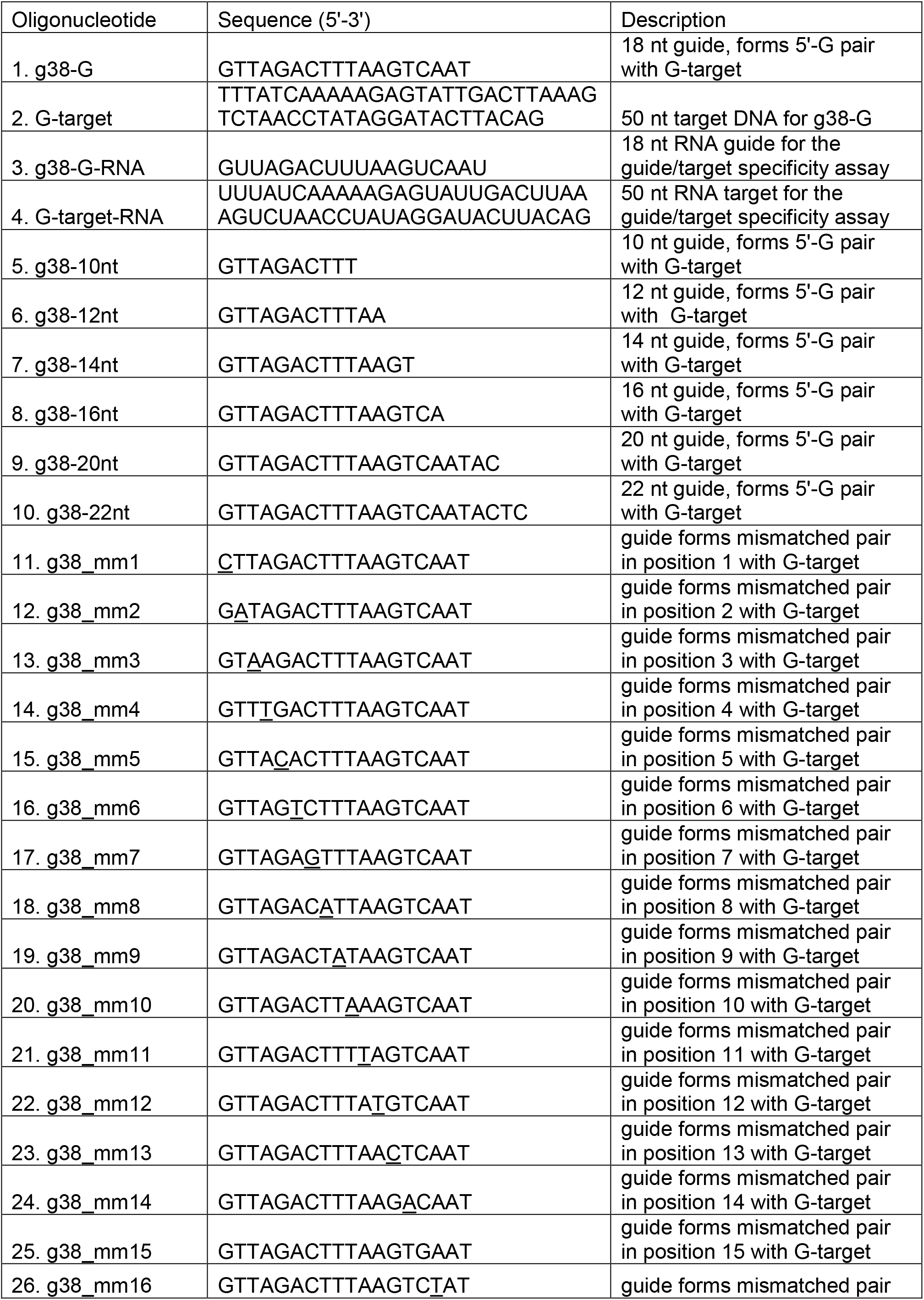
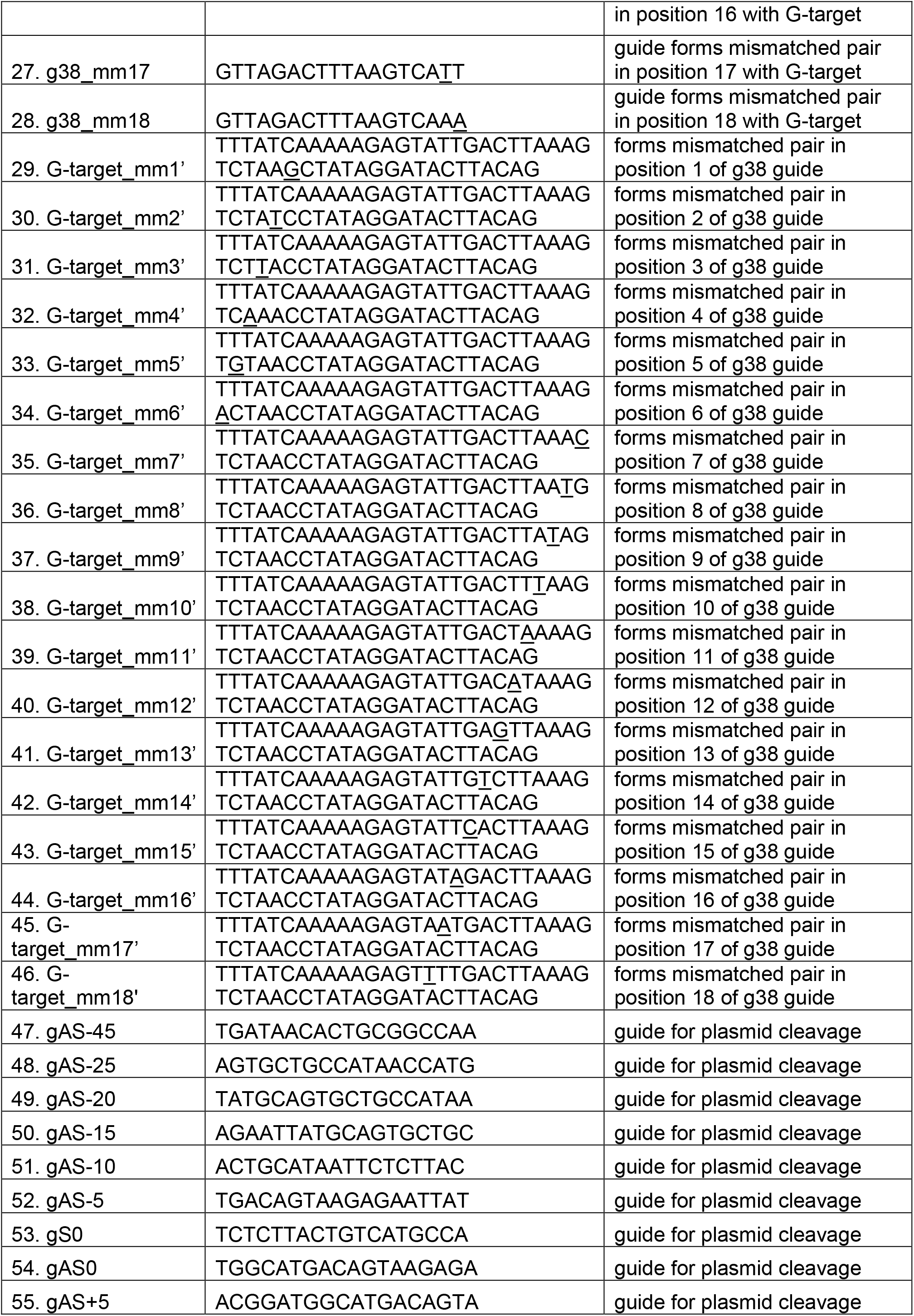
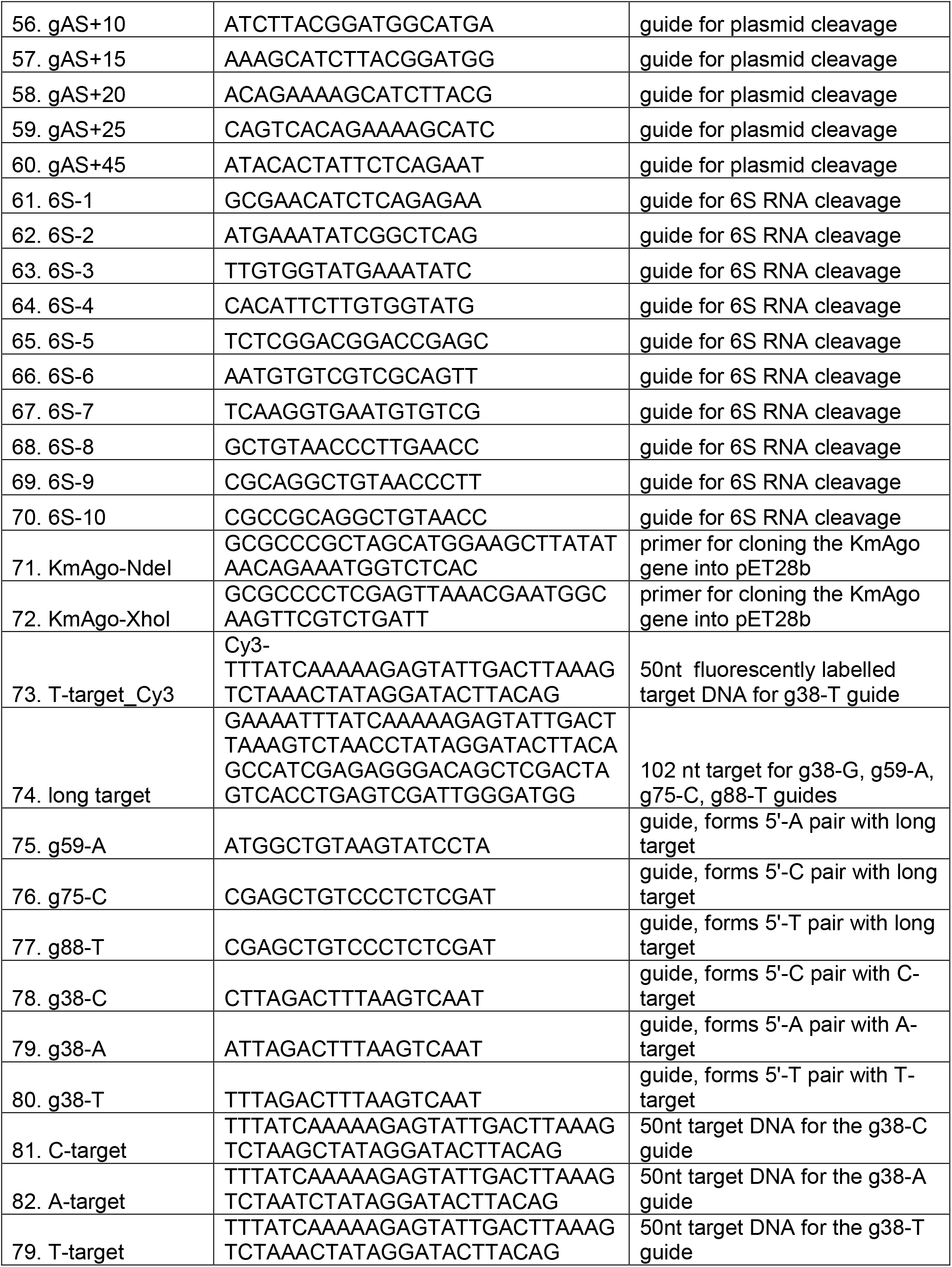
Sequences of oligonucleotides.

## References

1. Lisitskaya, L., Aravin, A.A. and Kulbachinskiy, A. (2018) DNA interference and beyond: structure and functions of prokaryotic Argonaute proteins. Nature communications, 9, 5165.

2. Makarova, K.S., Wolf, Y.I., van der Oost, J. and Koonin, E.V. (2009) Prokaryotic homologs of Argonaute proteins are predicted to function as key components of a novel system of defense against mobile genetic elements. Biology direct, 4, 29.

3. Swarts, D.C., Makarova, K., Wang, Y., Nakanishi, K., Ketting, R.F., Koonin, E.V., Patel, D.J. and van der Oost, J. (2014) The evolutionary journey of Argonaute proteins. Nature structural & molecular biology, 21, 743–753.

4. Willkomm, S., Makarova, K. and Grohmann, D. (2018) DNA-silencing by prokaryotic Argonaute proteins adds a new layer of defence against invading nucleic acids. FEMS microbiology reviews, 42, 376–387.

5. Ryazansky, S., Kulbachinskiy, A. and Aravin, A.A. (2018) The Expanded Universe of Prokaryotic Argonaute Proteins. mBio, 9, e01935–01918.

6. Olina, A.V., Kulbachinskiy, A.V., Aravin, A.A. and Esyunina, D.M. (2018) Argonaute Proteins and Mechanisms of RNA Interference in Eukaryotes and Prokaryotes. Biochemistry. Biokhimiia, 83, 483–497.

7. Swarts, D.C., Hegge, J.W., Hinojo, I., Shiimori, M., Ellis, M.A., Dumrongkulraksa, J., Terns, R.M., Terns, M.P. and van der Oost, J. (2015) Argonaute of the archaeon Pyrococcus furiosus is a DNA-guided nuclease that targets cognate DNA. Nucleic acids research, 43, 5120–5129.

8. Swarts, D.C., Jore, M.M., Westra, E.R., Zhu, Y., Janssen, J.H., Snijders, A.P., Wang, Y., Patel, D.J., Berenguer, J., Brouns, S.J.J. et al. (2014) DNA-guided DNA interference by a prokaryotic Argonaute. Nature, 507, 258–261.

9. Wang, Y., Juranek, S., Li, H., Sheng, G., Tuschl, T. and Patel, D.J. (2008) Structure of an argonaute silencing complex with a seed-containing guide DNA and target RNA duplex. Nature, 456, 921–926.

10. Wang, Y., Juranek, S., Li, H., Sheng, G., Wardle, G.S., Tuschl, T. and Patel, D.J. (2009) Nucleation, propagation and cleavage of target RNAs in Ago silencing complexes. Nature, 461, 754–761.

11. Willkomm, S., Oellig, C.A., Zander, A., Restle, T., Keegan, R., Grohmann, D. and Schneider, S. (2017) Structural and mechanistic insights into an archaeal DNA-guided Argonaute protein. Nature microbiology, 2, 17035.

12. Zander, A., Holzmeister, P., Klose, D., Tinnefeld, P. and Grohmann, D. (2014) Single-molecule FRET supports the two-state model of Argonaute action. RNA biology, 11, 45–56.

13. Zander, A., Willkomm, S., Ofer, S., van Wolferen, M., Egert, L., Buchmeier, S., Stockl, S., Tinnefeld, P., Schneider, S., Klingl, A. et al. (2017) Guide-independent DNA cleavage by archaeal Argonaute from Methanocaldococcus jannaschii. Nature microbiology, 2, 17034.

14. Hegge, J.W., Swarts, D.C., Chandradoss, S.D., Cui, T.J., Kneppers, J., Jinek, M., Joo, C. and van der Oost, J. (2019) DNA-guided DNA cleavage at moderate temperatures by Clostridium butyricum Argonaute. Nucleic acids research, 47, 5809–5821.

15. Kuzmenko, A., Yudin, D., Ryazansky, S., Kulbachinskiy, A. and Aravin, A.A. (2019) Programmable DNA cleavage by Ago nucleases from mesophilic bacteria Clostridium butyricum and Limnothrix rosea. Nucleic acids research, 47, 5822–5836.

16. Olina, A., Kuzmenko, A., Ninova, M., Aravin, A.A., Kulbachinskiy, A. and Esyunina, D. (2020) Genome-wide DNA sampling by Ago nuclease from the cyanobacterium Synechococcus elongatus. RNA biology, 1–12.

17. Kuzmenko, A., Oguienko, A., Esyunina, D., Yudin, D., Petrova, M., Kudinova, A., Maslova, O., Ninova, M., Ryazansky, S., Leach, D. et al. (2020) DNA targeting and interference by a bacterial Argonaute nuclease. Nature, 587, 632–637.

18. Lisitskaya, L., Petushkov, I., Esyunina, D., Aravin, A. and Kulbachinskiy, A. (2020) Recognition of double-stranded DNA by the Rhodobacter sphaeroides Argonaute protein. Biochemical and biophysical research communications, 533, 1484–1489.

19. Hunt, E.A., Evans, T.C., Jr. and Tanner, N.A. (2018) Single-stranded binding proteins and helicase enhance the activity of prokaryotic argonautes in vitro. PloS one, 13, e0203073.

20. Olovnikov, I., Chan, K., Sachidanandam, R., Newman, D.K. and Aravin, A.A. (2013) Bacterial argonaute samples the transcriptome to identify foreign DNA. Molecular cell, 51, 594–605.

21. Swarts, D.C., Szczepaniak, M., Sheng, G., Chandradoss, S.D., Zhu, Y., Timmers, E.M., Zhang, Y., Zhao, H., Lou, J., Wang, Y. et al. (2017) Autonomous Generation and Loading of DNA Guides by Bacterial Argonaute. Molecular cell, 65, 985–998.

22. Jolly, S.M., Gainetdinov, I., Jouravleva, K., Zhang, H., Strittmatter, L., Bailey, S.M., Hendricks, G.M., Dhabaria, A., Ueberheide, B. and Zamore, P.D. (2020) Thermus thermophilus Argonaute Functions in the Completion of DNA Replication. Cell, 182, 1545–1559 e1518.

23. Kaya, E., Doxzen, K.W., Knoll, K.R., Wilson, R.C., Strutt, S.C., Kranzusch, P.J. and Doudna, J.A. (2016) A bacterial Argonaute with noncanonical guide RNA specificity. Proceedings of the National Academy of Sciences of the United States of America, 113, 4057–4062.

24. Sheng, G., Zhao, H., Wang, J., Rao, Y., Tian, W., Swarts, D.C., van der Oost, J., Patel, D.J. and Wang, Y. (2014) Structure-based cleavage mechanism of Thermus thermophilus Argonaute DNA guide strand-mediated DNA target cleavage. Proceedings of the National Academy of Sciences of the United States of America, 111, 652–657.

25. Elkayam, E., Kuhn, C.D., Tocilj, A., Haase, A.D., Greene, E.M., Hannon, G.J. and Joshua-Tor, L. (2012) The structure of human argonaute-2 in complex with miR-20a. Cell, 150, 100–110.

26. Wassarman, K.M. (2018) 6S RNA, a Global Regulator of Transcription. Microbiology spectrum, 6.

27. Liu, Y., Esyunina, D., Olovnikov, I., Teplova, M., Kulbachinskiy, A., Aravin, A.A. and Patel, D.J. (2018) Accommodation of helical imperfections in *Rhodobacter sphaeroides* Argonaute ternary complexes with guide RNA and target DNA. Cell reports, 24, 453–462.

28. Song, J., Hegge, J.W., Mauk, M.G., Chen, J., Till, J.E., Bhagwat, N., Azink, L.T., Peng, J., Sen, M., Mays, J. et al. (2020) Highly specific enrichment of rare nucleic acid fractions using Thermus thermophilus argonaute with applications in cancer diagnostics. Nucleic acids research, 48, e19.

29. He, R., Wang, L., Wang, F., Li, W., Liu, Y., Li, A., Wang, Y., Mao, W., Zhai, C. and Ma, L. (2019) Pyrococcus furiosus Argonaute-mediated nucleic acid detection. Chemical communications, 55, 13219–13222.

30. Lapinaite, A., Doudna, J.A. and Cate, J.H.D. (2018) Programmable RNA recognition using a CRISPR-associated Argonaute. Proceedings of the National Academy of Sciences of the United States of America, 115, 3368–3373.

31. Dayeh, D.M., Cantara, W.A., Kitzrow, J.P., Musier-Forsyth, K. and Nakanishi, K. (2018) Argonaute-based programmable RNase as a tool for cleavage of highly-structured RNA. Nucleic acids research, 46, e98.

32. Shin, S., Jung, Y., Uhm, H., Song, M., Son, S., Goo, J., Jeong, C., Song, J.J., Kim, V.N. and Hohng, S. (2020) Quantification of purified endogenous miRNAs with high sensitivity and specificity. Nature communications, 11, 6033.

